# The local topological free energy of proteins

**DOI:** 10.1101/2021.01.06.425494

**Authors:** Quenisha Baldwin, Eleni Panagiotou

**Affiliations:** Department of Biology, Tuskegee University, AL 36088, USA < >; Department of Mathematics and SimCenter, University of Tennessee at Chattanooga, TN 37403, USA < >

**Keywords:** Protein structure, protein folding, topology, Writhe, Torsion

## Abstract

Protein folding, the process by which proteins attain a 3-dimensional conformation necessary for their function, remains an important unsolved problem in biology. A major gap in our understanding is how local properties of proteins relate to their global properties. In this manuscript, we use the Writhe and Torsion to introduce a new local topological/geometrical free energy that can be associated to 4 consecutive residues along protein backbone. By analyzing a culled protein dataset from the PDB, our results show that high local topological free energy conformations are independent of sequence and may be involved in the rate limiting step in protein folding. By analyzing a set of 2-state single domain proteins, we find that the total local topological free energy of these proteins correlates with the experimentally observed folding rates reported in [29].

## 1 Introduction

Protein folding is the process by which a protein attains a unique three-dimensional conformation necessary for its function [2]. Many different models of protein folding have been proposed, all of which aim to understand the free energy barrier associated with the transition from the unfolded configuration to the native state of the protein [1,3,4,9–12,14–18,20,22–24,31–33]. To describe this process, which involves a myriad of length scales, it is necessary to have a meaningful characterization of the 3-dimensional configuration of proteins across length scales. In this manuscript we introduce characterizations of protein conformations using tools from mathematics, related to knot theory, that apply to all protein length scales. In particular, we focus at characterizing the local conformations proteins (those of 4 consecutive CA atoms) and exploring the relation to the global configuration of the protein and protein kinetics.

Folded proteins are defined by their primary, secondary, tertiary and quaternary structure [2]. The primary structure refers to the protein amino acid sequence. The secondary structure refers to a sequence of 3-dimensional building blocks the protein attains (beta sheets, alpha helices, coils). The tertiary structure refers to the 3-dimensional conformation of the entire polypeptide chain. The quaternary structure of a protein comprises of 2 or more polypeptide chains. More refined methods to characterize protein conformations than these classifications are also used. For example, at the level of amino acids, the Ramachandran plot, is a traditional way to capture the geometrical signatures of amino acids in terms of their dihedral angles in 3-space. At the length scale of the entire protein, the number of sequence-distant contacts is a way to describe the conformation of the protein, which has shown a remarkable correlation with experimentally observed folding rates [19, 20, 27, 28].

In the last decade, measures from knot theory have been applied to proteins to classify their conformations [5–7, 25, 34]. One of the simplest measures of conformational complexity of proteins that does not require an approximation of the protein by a knot dates back to Gauss; the Writhe of a curve. Studies have applied the Gauss linking integral to measure the entanglement of the protein backbone by taking the entire backbone of the protein or by looking at linking between parts of the protein. Both approaches have found a correlation between folding rates and these measures of conformational complexity [5–7, 25]. However, this does not answer how local properties of proteins relate to its tertiary structure. The protein backbone, represented by its CA atoms, can attain interesting conformations even with as little as 4 residues. To our knowledge no study has focused at exploring the local topology/geometry of the proteins at the length scale of 4 using the Gauss linking integral and Torsion. In this manuscript, we use the Writhe to define a novel topological/geometrical free energy that can be assigned locally to the protein. We do the same also using the Torsion. We use this free energy to identify high local topological free energy conformations. Our results show that the high local topological free energy conformations are independent of the local sequence and may be involved in the rate limiting step in protein folding. By analyzing a set of 2-state, single domain proteins, we show that the previously reported experimental folding rates in [29] correlate with the total local topological free energy of the proteins, with slower folding rates associated to higher total local topological/geometrical free energy.

The paper is organized as follows: Section 2 describes the topological and geometrical functions for characterizing 3-dimensional conformations used in this paper. Section 3 describes our results. Finally, in Section 4, we summarize the findings of our analysis.

## 2 The local topological/geometrical free energy of proteins

In Section 2.1 we give the definition of the mathematical tools that we will use in this manuscript. In Section 2.2 we introduce the novel definition of a local topological free energy of the protein backbone crystal structure.

### 2.1 Measures of topological/geometrical complexity

We represent proteins by their CA atoms, as linear polygonal curves in space. A measure of conformational complexity of curves in 3-space is the Gauss linking integral. When applied to one curve, this integral is called the Writhe of a curve:

#### Definition 2.1.

(Writhe). For a curve *ℓ* with arc-length parameterization *γ*(*t*), the Writhe, *Wr*, is the double integral over *l*:

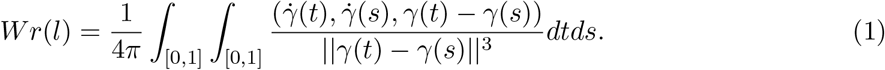

The Writhe measures the average algebraic sum of crossings of the projection of the curve with itself over all possible projection directions. It is a measure of the number of times a chain winds around itself and can have both positive and negative values.

The total Torsion of the chain, describes how much it deviates from being planar and is defined as:

#### Definition 2.2.

The *Torsion* of a curve *ℓ* with arc-length parameterization *γ*(*t*) is the double integral over *l*:

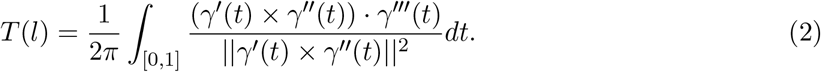

The Writhe and the Torsion have successfully been applied to study entanglement in biopolymers and proteins in particular [5–7, 21, 25, 30].

An important property of the Gauss linking integral and the Torsion which makes them useful in practice is that they can be applied to polygonal curves of any length to characterize 3-dimensional conformations at different length scales. In this work, we use the Writhe and the Torsion to characterize local 3-dimensional conformations of a protein at the length scale of 4 residues, we call this the *local Writhe* and the *local Torsion*, respectively.

#### Definition 2.3.

We define the *local Writhe of a residue* (resp. *local Torsion*), represented by the CA atom *i* to be the Writhe (resp. *Torsion*) of the protein backbone connecting the CA atoms *i, i* + 1, *i* + 2, *i* + 3.

The local Writhe is a measure of the local orientation of a polygonal curve and a measure of its compactness. For example a very tight right-handed turn (resp. left-handed) will have a positive (resp. negative) Writhe value close to 0.5 (resp. −0.5), while a relatively straight segment will have a value close to 0. Similarly, the Torsion is 0 for a planar segment and increases to ±1 as the segment deviates from being planar. It is possible to have low Writhe and high Torsion and the opposite. Figure 1 shows examples of the Writhe and Torsion values when applied globally to the entire protein or locally, to 4 consecutive residues of the protein.

**Figure 1:**
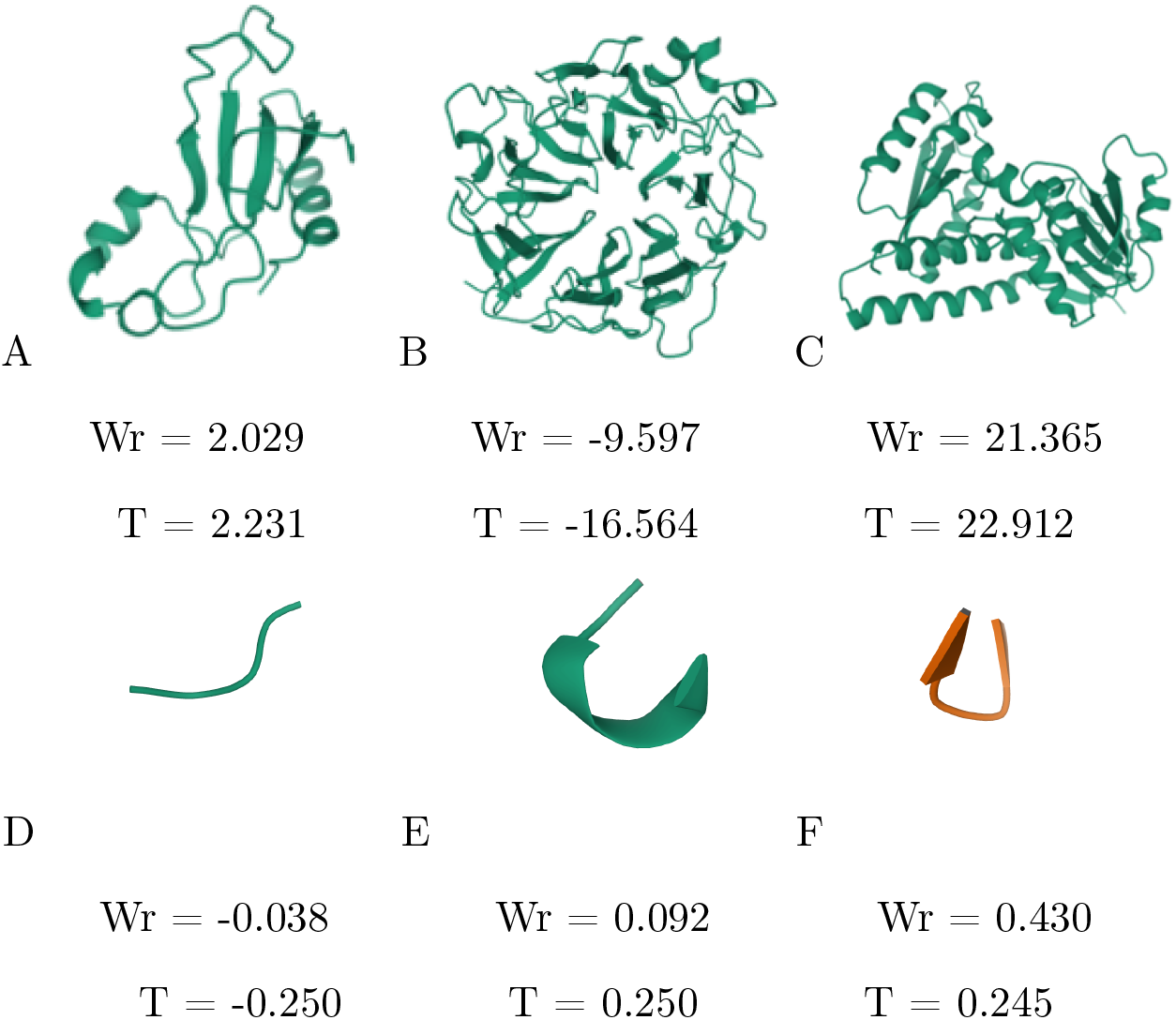
Example of global vs. local Writhe (Torsion) (A) Global Writhe and Torsion of PDB: 1A2P (resp. (B) 1A12, (C) 1A4I), are shown to increase in absolute value as length and complexity of protein increases. (D) Local Writhe and Torsion values of PDB: 16PK residues 1-4 (E) residues 4-8 and (F) 1GK9 residues 92-96 shown.

### 2.2 Topological/Geometrical free energy

To assign a local topological/geometrical free energy along a protein backbone, we use a method inspired by the framework used in [26] for identifying exotic geometries of hydrogen bonds derived through DFT calculations. We first derive the distributions of the local Writhe and local Torsion in the ensemble of folded proteins. Then for each local conformation of a given protein we compare its local Writhe (resp. Torsion) value to those of the ensemble and a free energy is assigned to it based on the population of that value in the ensemble. We can do the same for the global topology/geometry of the entire protein.

Let *X* denote a topological parameter (such as Writhe or Torsion). We compute the distribution of X in the folded state ensemble. To do this in practice, we use a culled subset of the crystal structures provided in the PDB. Namely, we use the dataset of unbiased, high-quality 3-dimensional structures with less than 60% homology identity from [35].

Let *d_X_* denote the density (ie. the number of occurrences) of X in the folded ensemble (*d_Wr_*, *d_T_*, respectively). Let *m_X_* (resp. *m_Wr_*, *m_T_*) denote the maximum occurrence value for X. To any value *p* of X, we associate a normalized quantity, which we will call, *topological free energy*:

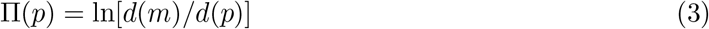

We denote Π_*Wr*_, Π_*T*_ the topological free energy in Writhe and Torsion, respectively.

We note that the above definition of topological free energy can be applied to different lengths of the protein, by measuring the parameter X for *n* consecutive residues at a time. In this manuscript, we will focus at n = 4 which is the smallest possible n that can be used to define Writhe and Torsion. We will say that a residue has a *high local topological free energy in a parameter X* (or is *rare in a parameter X*) (where *X* is local Writhe or local Torsion) if its value *X* = *p* is such that Π(*p*) ≥ *w*, where w is a threshold corresponding to the 95^th^ percentile of Π-values across the set of folded proteins. We stress that our definition of rare residue involves 4 consecutive residues, starting from the one we identify as rare.

We note that the free energy that we defined captures topological effects in long chains. For short sequences of 4 residues, the topology is trivial, but the geometry is not. Nevertheless, since we use the Writhe, defined through the Gauss linking integral, a conventional tool in topology, we will use the term topological free energy at all length scales, even at 4 residues.

## 3 Results

In Section 3.1 we present our results on the local topology/geometry of the culled PDB ensemble. In Section 3.2 we examine local conformations of high topological/geometrical free energy. In Section 3.3 we analyze a set of 2-state singe domain proteins to examine the relation between the total local topological free energy along the protein backbone and the experimentally observed folding rate of the protein.

### 3.1 Local topology in the PDB

In this section we present the analysis of the local topology/geometry of the culled ensemble of the PDB proteins at 4 consecutive residues at a time along the entire backbone.

Figures 2A and B show the local Writhe and local Torsion distributions in the PDB culled ensemble, respectively. The local Writhe and local Torsion show one local maximum at positive values and one at negative. This suggests a well defined pattern in the local conformation of folded proteins. A peak at a positive Writhe or Torsion value suggests high presence of right-handed local conformations. A peak at a negative Writhe or Torsion value suggests high presence of left-handed local conformations. This could be a manifestation of the secondary structure of the proteins analyzed: Namely, 98% of the PDB protein culled sample contain at least one helix which contribute positive writhe values and 91% of the PDB protein culled sample contain at least one beta sheet, and *β*-strands may contribute small negative Writhe values [25]. The local Writhe values are concentrated between −0.2 and 0.2. The local Writhe maxima occur at approximately −0.01 and 0.8. Interestingly, the local Torsion distribution seems to be almost entirely concentrated in two values, a positive and a negative, almost equally populated each. The local Torsion shows local maxima at −0.25 and 0.25. Despite this general similarity, the distributions of Writhe and Torsion are different. Both distributions of local Writhe and Torsion have more pronounced peaks than the global Writhe and Torsion distributions discussed in the Appendix. This may indicate a strong pattern in the local configuration of proteins which influences the global conformation of the proteins.

**Figure 2:**
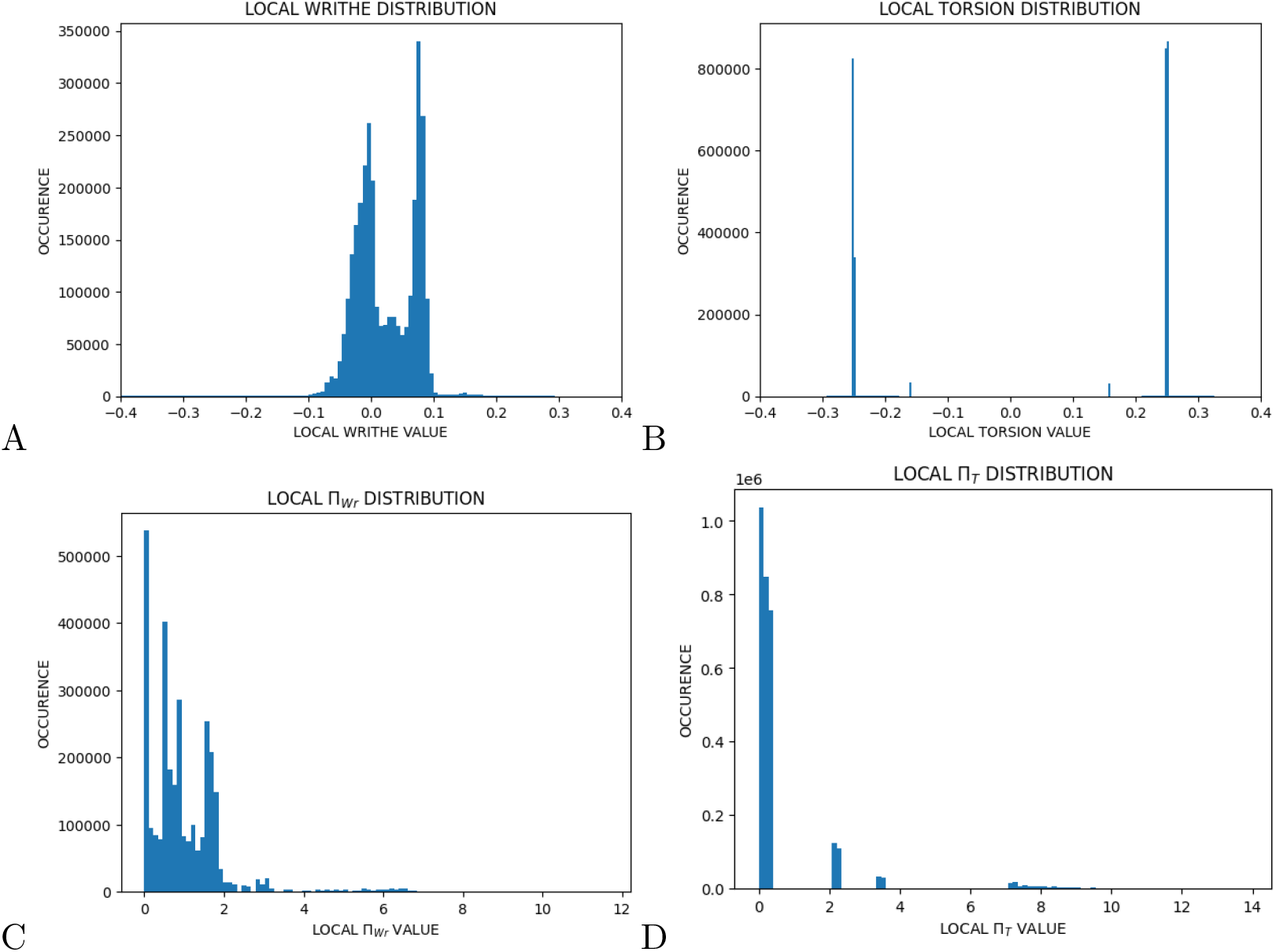
Distribution of the local topology in the PDB. (A) The local Writhe. (B) The local Torsion. Both distributions are bimodal but very different from each other and from the global Writhe and Torsion distributions shown in the Appendix. (C, D) The local Π distribution for Writhe and Torsion, respectively.

Figures 2C and D show the Π_*Wr*_-values and Π_*T*_-values, respectively. We see that most Π values are less than 2, indicative of low topological free energy. It is interesting to point out that these distributions are broader than those of the Π values of the global Writhe and Torsion (not shown here). This suggests that there may be low global topological free energy configurations, which contain high local topological free energy configurations.

### 3.2 High local topological free energy conformations in proteins

In this Section we will focus on those conformations in the 95^th^ percentile of the distributions, which correspond to high Π values, which we associate with high local topological/geometrical free energy. For simplicity, we will also refer to them as rare conformations. The 95^th^ percentile of the Π_*Wr*_-value distribution in the PDB corresponds approximately to absolute Writhe values greater than 0.1. The 95^th^ percentile of the Π_*T*_-value distribution in the PDB corresponds approximately to absolute Torsion values greater than 0.3 or smaller than 0.1. Thus, rare local Writhe values correspond to high Writhe in absolute value, while rare local Torsion values may be values close to 0 in absolute value. This suggests that rare conformations in Writhe may not necessarily be rare in Torsion and the opposite. In general, the high local Writhe values could correspond to tight right-handed turns and the low local Torsion values could correspond to almost planar conformations.

In Section 3.2.1 and Section 3.2.2, respectively, we examine if high local topological free energy conformations are related to secondary structure elements and/or specific amino acid types.

#### 3.2.1 High local topological free energy conformations and secondary structure in the PDB

To better understand the meaning of values of local Writhe and Torsion at the 95^th^ percentile of the Π-value distribution, we examine the correlation between these values and secondary structure elements.

Our analysis showed that the entire local conformation lies within the secondary structure of its first residue. Figure 3 shows the distribution of the first residue rare local configurations in secondary structure elements. We see that 41% of the rare conformations in both local Writhe and Torsion are in coils, 40% in helices and 19% in *β* sheets. We first notice that conformations with high local topological free energy are more frequent in coils and helices. The high percentage of high local topological free energy conformations in coils is in agreement with our intuition that these may be associated to high free energy. It may be surprising on the other hand that a larger percentage in total of the high local topological free energy conformations are in beta sheets and helices, which are stable secondary structures. We stress that even if the locations of rare conformations in Writhe and Torsion are similarly distributed across secondary structures, they are not pointing to the same residues.

**Figure 3:**
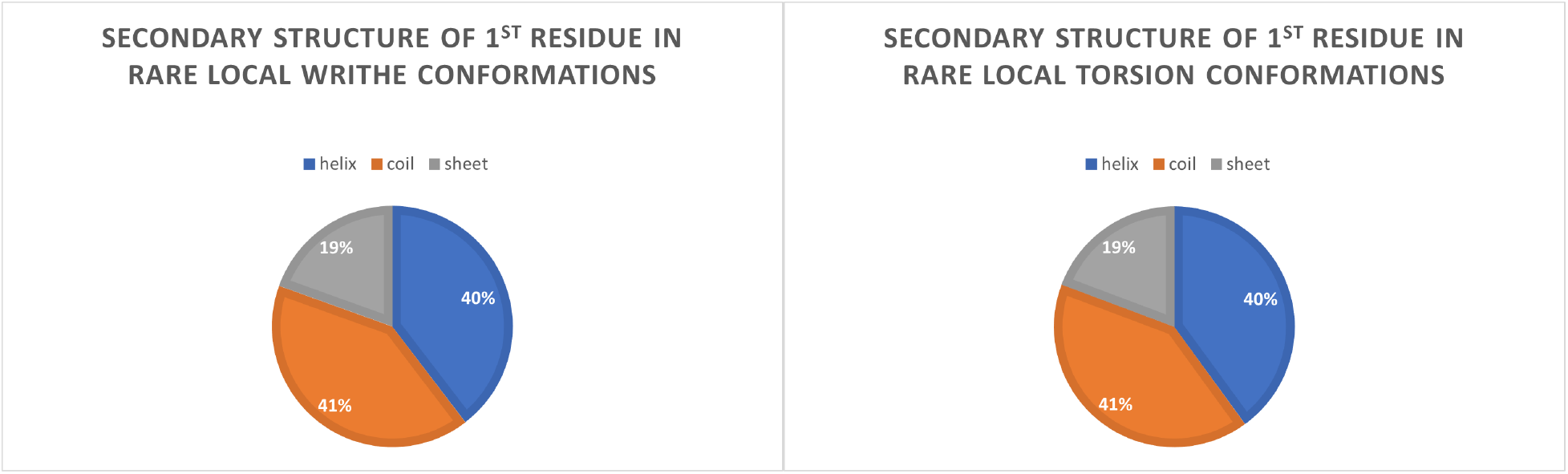
The distribution in secondary structure elements of the first residue in high local topological/geometrical free energy configurations in Writhe (Left) and in Torsion (Right) in the PDB culled data set.

#### 3.2.2 High local topological free energy conformations and amino acid type in the PDB

Amino acids have preferred dihedral angle distributions, specific sizes and other amino acid type dependent physical properties. It is natural therefore to examine whether there is a correlation between a residue being part of a rare conformation and its amino acid type.

Figure 4 shows the frequency of each amino acid in the PDB culled dataset versus the frequency each amino acid appears as part of a high local topological free energy conformation. Overall, we see that the frequency by which an amino acid occurs in a rare conformation is the same as the frequency by which it appears in the PDB culled dataset. This suggests that high local topological free energy configurations are independent of their sequence. Exceptions might be phenylalanine and histidine in both local Writhe and Torsion. Phenylalanine appears to be favoring rare local conformations while histidine is not favored in rare local conformations.

**Figure 4:**
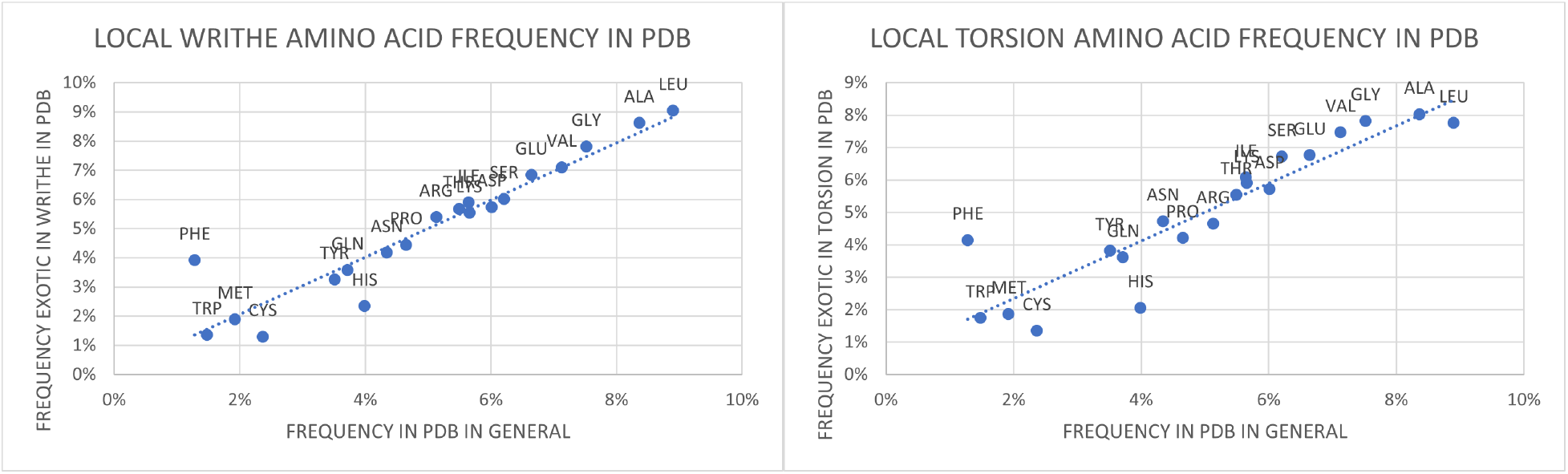
Frequency of amino acid types in high local topological free energy configurations in Writhe (Left) and in Torsion (Right) are plotted with frequency of amino acid types in the PDB in general. We find a linear fit with R^2^ = 0.8297 (resp. R^2^ = 0.885) suggesting that high local topological free energy configurations are not related to local amino acid sequence.

We next examine the handedness of the exotic configurations led by each amino acid. As a proxy for handedness we simply use the sign of the Writhe and torsion, where positive sign indicates right-handed, while negative sign indicates left-handed. We find that the percentage of positive Writhe values for each amino acid fall within the range of 56-65% (with the exception of Methionine which is 69% positive in local Writhe and cysteine which is 49% positive in local Writhe). These results suggest a small but consistent preference for positive local Writhe and Torsion values for high local topological free energy conformations, representative of right-handed conformations.

We also examine the average absolute local Writhe and Torsion for each amino acid in a high local topological free energy conformation, shown in Figure 5. The local Writhe varies roughly between 0.01 and 0.03 and the local Torsion varies roughly from 0.025 to 0.07. Methionine has the highest average absolute local Writhe value. Methionine is hydrophobic and uncharged, it is, therefore, possible that methionine is part of a turning conformation in order to tuck away from the hydrophilic extracellular space. Amino acids glutamic acid, tyrosine, isoleucine and alanine are among those amino acids that lead conformations with the highest average absolute local torsion, which represent the most non-planar conformations.

**Figure 5:**
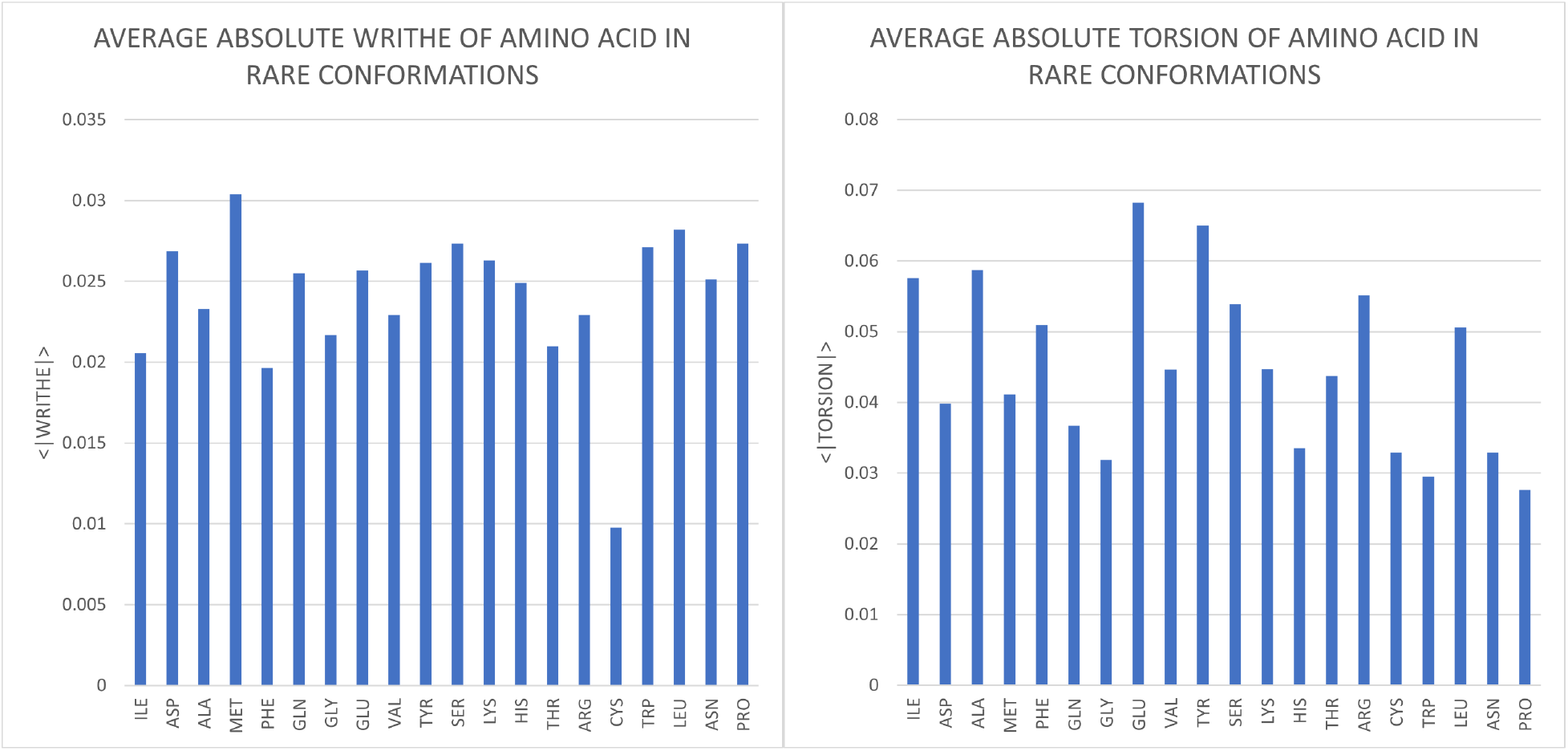
Average absolute Writhe values (Left) and Torsion values (Rights) of high local topological free energy conformations containing for each amino acid they contain.

Figure 6 shows the average local topological free energy for local Writhe and Torsion for each amino acid when involved in a high topological free energy configuration. Our results show that CYS has the lowest Π_*Wr*_-values, but one of the highest Π_*T*_-values, while PHE has one of the highest Π_*Wr*_-values and one of the smallest Π_*T*_-values.

**Figure 6:**
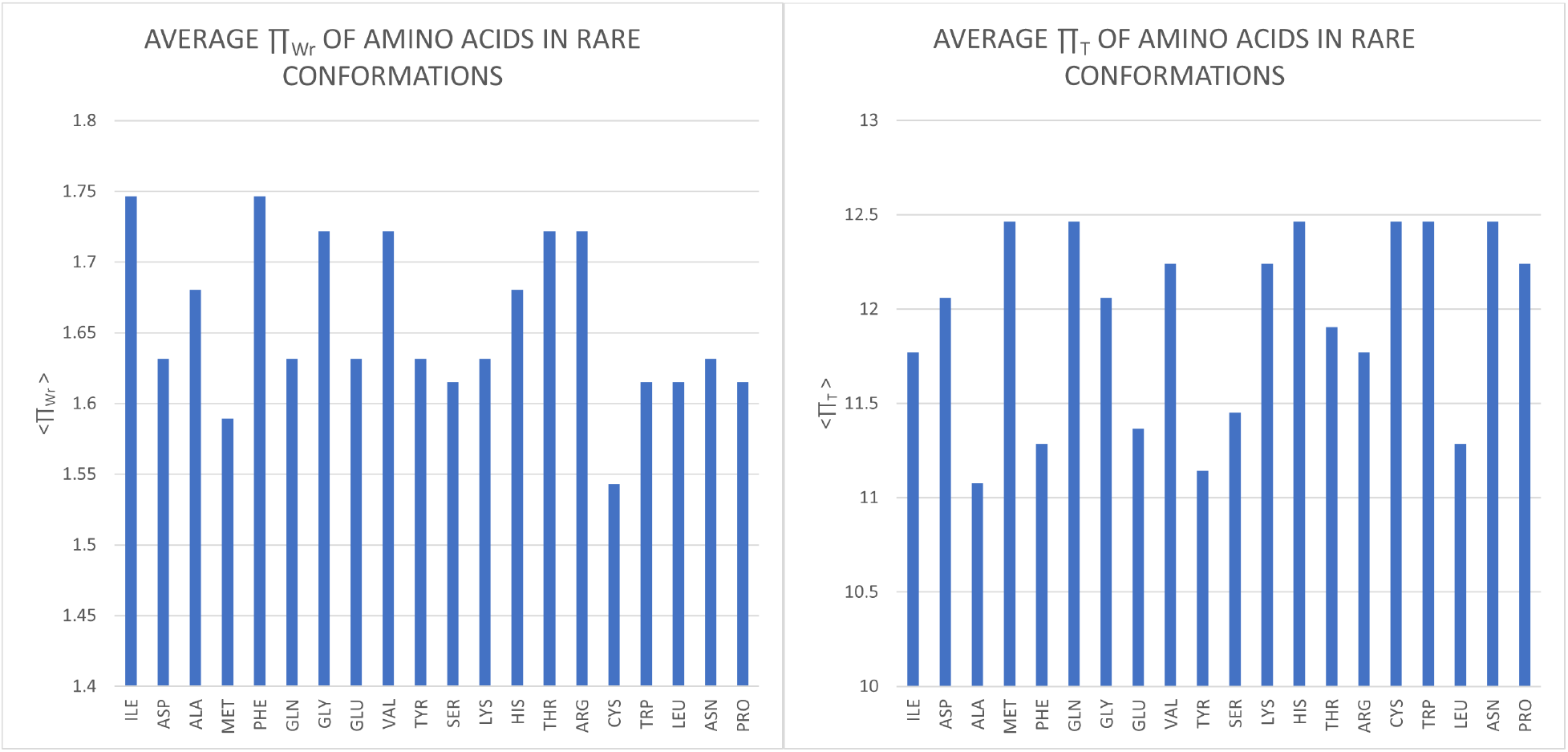
Average Π_Wr_ (Left) and Π_T_ (Right) values for high local topological free energy conformations and the amino acids they contain.

### 3.3 Local topological free energy and protein folding rates

In the previous paragraph we found no significant amino acid preference for being in a high local topological free energy conformation. We may thus infer that the rare local conformations are not related to the local protein sequence. In this section we will examine how the local conformation of proteins may be related to protein folding kinetics.

We analyze a set of simple, single domain, non-disulfide-bonded proteins that have been reported to fold in a concerted, all-or-none, two-state fashion, whose experimental folding rates in water were obtained in [29]. In [25] it was shown that the logarithm of the experimental folding rate decreases with decreasing global Writhe and Torsion of the protein backbone. In this section we examine how the total local topological free energy correlates with folding rate.

First we compute the total local free energy by adding the Π values along the backbone of a protein. Figure 7 shows the logarithm of the experimental folding rate versus the total local topological free energy in Torsion (total sum of Π_*T*_ values along the backbone). Our results show that the folding rate decreases with increasing total local topological free energy in Torsion along the entire backbone (with R^2^ = 0.38). Note that the sum of Π_*T*_ values increases with the more “rare” a local conformation is. Interestingly, we do not find a correlation between the folding rate and the total local Π_*Wr*_-values for this dataset.

**Figure 7:**
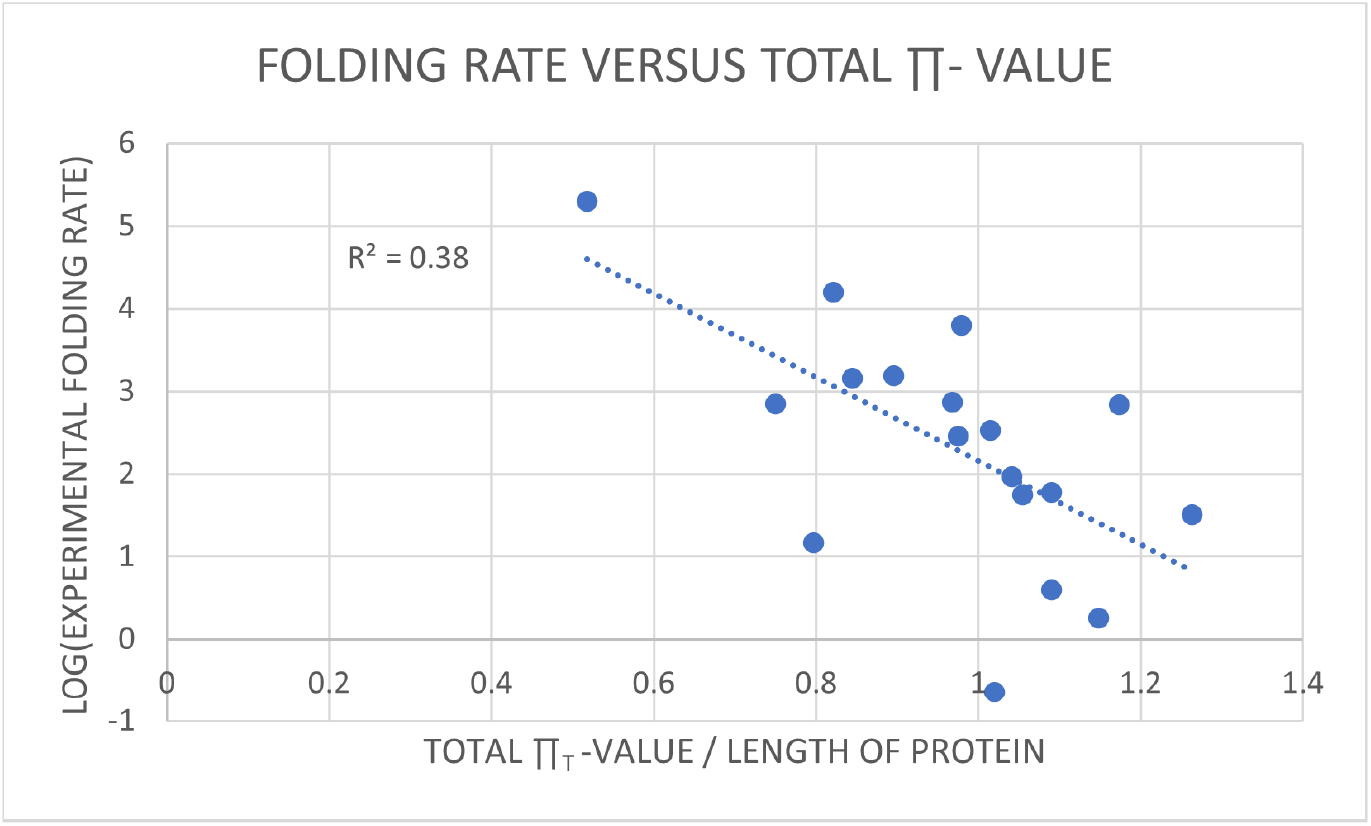
The logarithm of the experimentally observed folding rate of a set of 2-state proteins as a function of the normalized sum of local topological free energy in Torsion (the sum of Π_T_-values along the protein backbone).

We also examine the logarithm of the experimental folding rate versus the number of rare (high local topological free energy) residues a protein has in Figure 8. Interestingly, we find that overall the folding rate decreases with increasing number of high local topological free energy in Writhe local conformations, which suggests that these rare local conformations may represent global energy barriers in the protein.

It is interesting to point out that this trend is apparent only in Writhe and not in Torsion, even though the opposite was true for the total local topological free energy.

**Figure 8:**
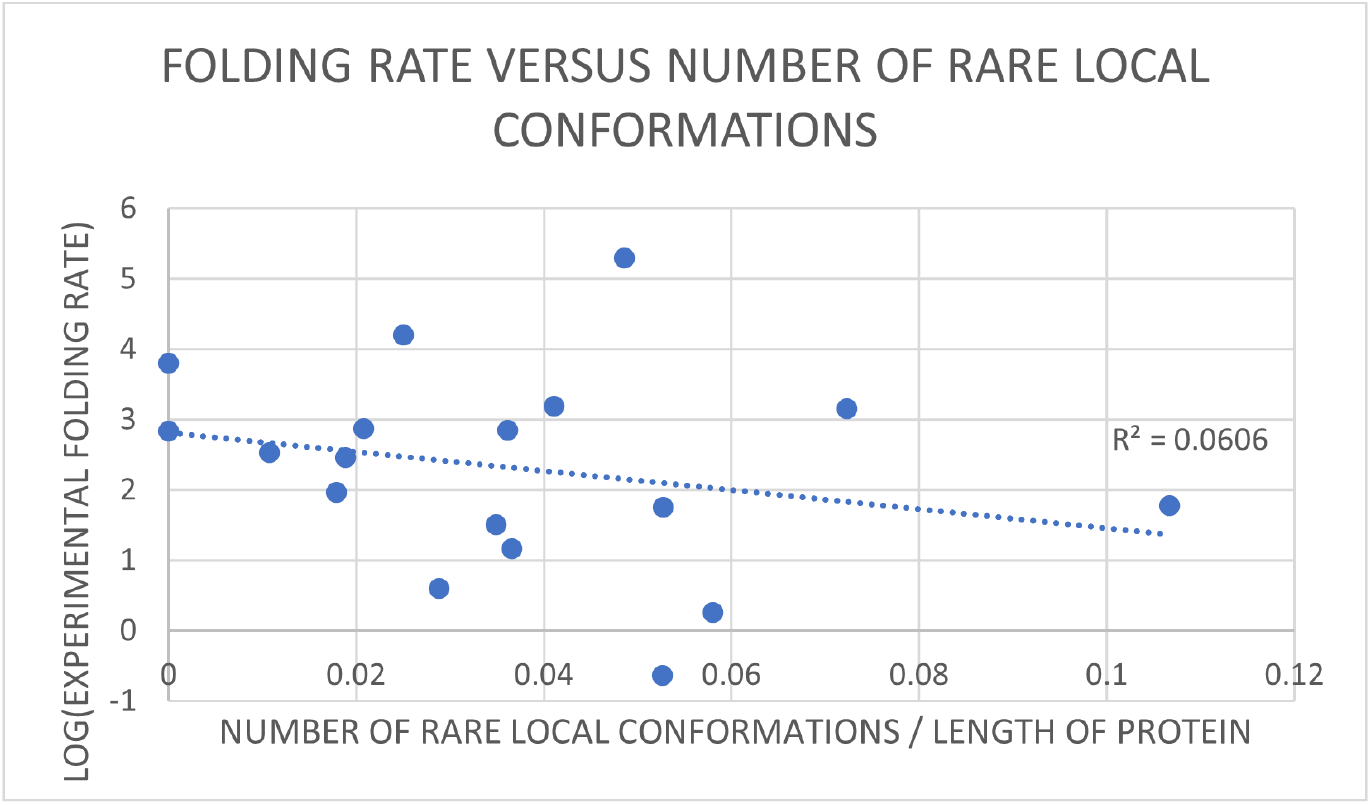
The logarithm of the experimentally observed folding rate of a set of 2-state proteins as a function of the normalized number of high local topological free energy conformations in Writhe.

### 3.4 Local topological free energy and *ϕ* values

The effect of amino acid sequence on the tertiary structure of a protein is studied experimentally though chemical scannings. Chemical scanning consists in substituting amino acids along a protein backbone (for a protein known to fold) and exploring its folding after the substitution to its native state. This comparison is done using a quantity called *ϕ* value, which reflects how much the mutated amino acid is involved in the key contacts established during the folding process. A *ϕ* value that is equal to 0, suggests that the mutation has no effect on the structure and that the region surrounding the mutation is unfolded in the transition state. A *ϕ* value that is equal to 1 means that the local structure around the mutation closely resembles the structure of the native state. Therefore, the *ϕ* value represents how a specific amino acid in the sequence has an effect in the global structure of the protein, *ϕ* =1 assuming not important at the rate limiting step. Here we focus on exploring whether there is a relation between Π-values and experimentally observed *ϕ* values for a set of well studied proteins: barnase, FK506 binding protein (FKBP12), chrymotypsin inhibitor (CI2) and src SH3 domain (SH3).

We calculate the Π_*Wr*_ values along the backbone of the proteins and compare them to the experimentally reported *ϕ* values along their backbone [8,13]. Our results, shown in Figure 9, display an overall decrease in *ϕ* as Π_*Wr*_ increases. This further supports our findings showing that high Π_*Wr*_ values are associated with high free energy conformations which are sensitive in the rate-limiting step of the folding process. This is not found in this data for Torsion.

**Figure 9:**
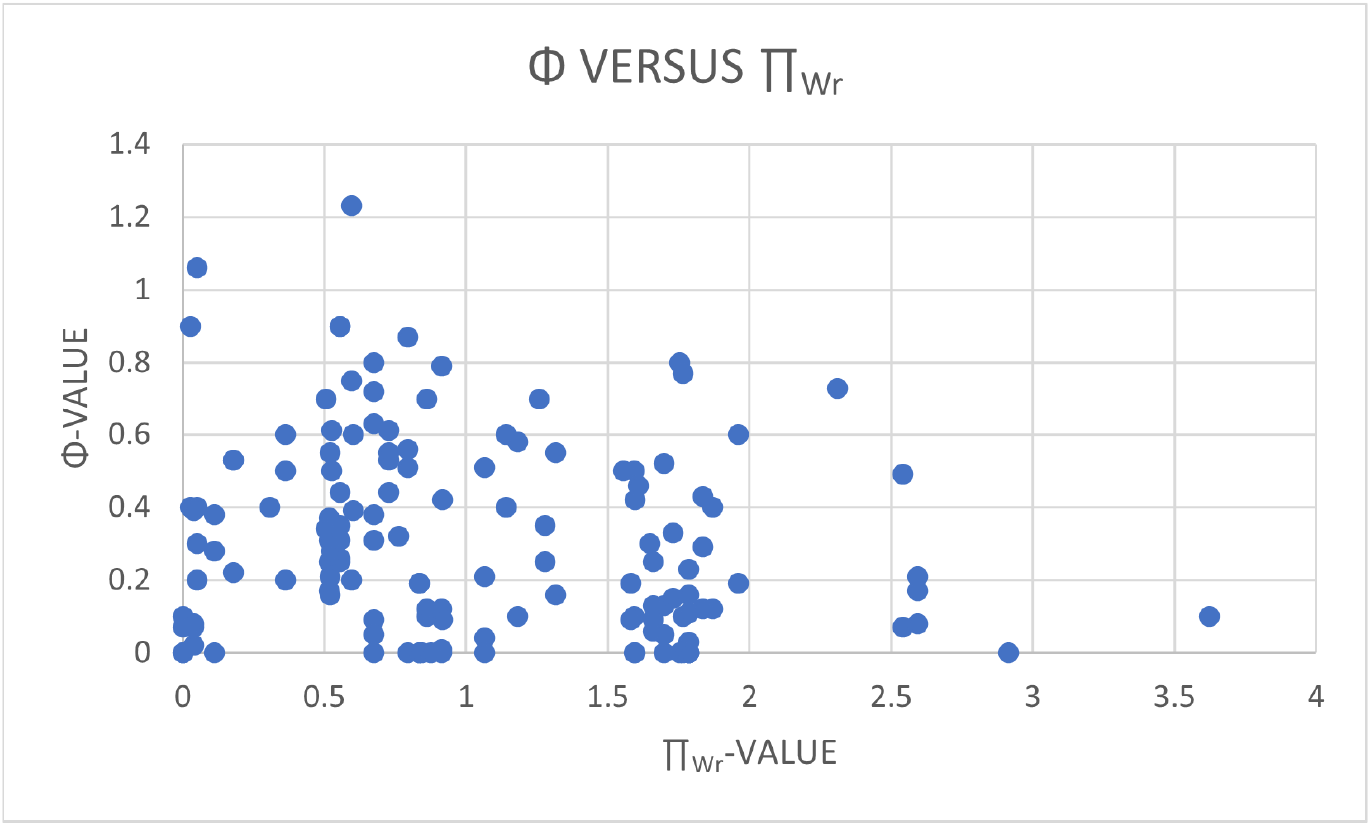
ϕ-values versus the Π-values of proteins barnase (PDB ID: 1BRS), FKBP12 (PDB ID: 1FJK), CI2 (2CI2) and SH3 (PBD ID: 1SRL).

## 4 Discussion

We used the local topology/geometry of protein crystal structures alone to associate a novel local topological/geometrical free energy to the protein backbone amino acids. By using a culled protein data set from the PDB we derived the distributions of the local Writhe and local Torsion values. Using these, we computed a local topological free energy for each protein. For the PDB culled data set we studied in this manuscript, we found that the high local topological free energy conformations in Writhe and in Torsion are preferably in coils and helices. Our results showed that there is no correlation between local sequence and detecting high local topological free energy conformations. Interestingly, our results suggest that these high local topological free energy conformations are related to the global conformations of the proteins. Namely, by focusing on a well studied set of 2-state proteins, we found that the logarithm of experimental folding rates is decreasing with the total local topological free energy in Torsion along the protein backbone. We also found an overall decrease of the logarithm of the folding rate with the number of high local topological free energy conformations in Writhe per protein. Our results also showed that *ϕ* values decrease with increasing local topological free energy in Writhe. These results point to the fact that the local topological free energy in Torsion and in Writhe capture different information about proteins. Namely, the total local topological free energy in Torsion better captures a free energy for the entire protein backbone, while the local topological free energy in Writhe can be better used to identify local conformations which capture important features of the entire protein conformation.

## 6 Acknowledgments

We thank Dr. Bobby Sumpter and Dr. Cristian Micheletti for very helpful discussions.

## 7 Appendix: The global topology of proteins in the PDB

In this section we analyze the three-dimensional configuration of the entire protein backbone of the PDB culled protein dataset.

The normalized values of the Writhe by the length of the proteins is shown in Figure 10(A). We see a bimodal distribution with a peak at positive values of Writhe and a smaller one at negative Writhe values. The distribution is overall skewed to the positive values of Writhe, ranging from approximately −0.2 to 0.6. The skewness of the distribution to positive Writhe values indicates a preference for right-handed conformations in the proteins. This could be a manifestation of the secondary structure of the proteins analyzed or of the local Writhe values, discussed in Section 3.1. However, we note that even though the secondary structure element Writhe values add to the global Writhe of the protein, there can be helical proteins with negative Writhe and proteins with no helices that have positive Writhe.

**Figure 10:**
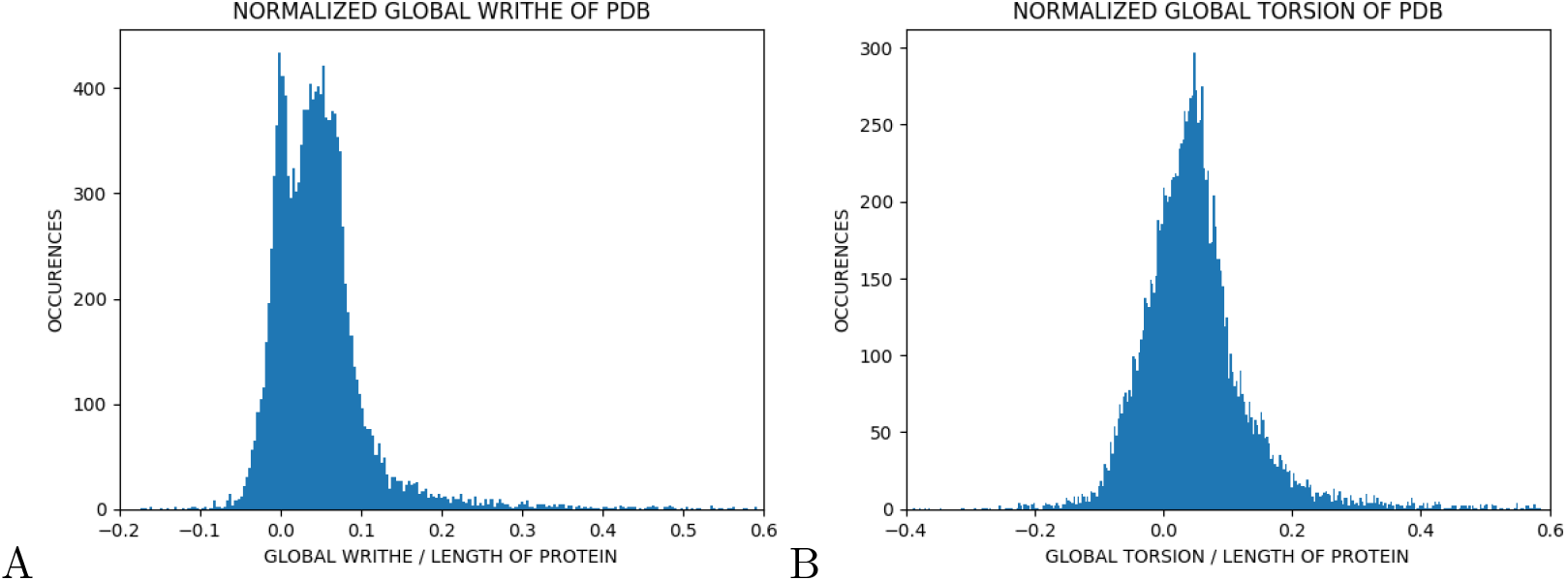
Distribution of the global topology/geometry of the PDB culled dataset.(A) Writhe of a protein normalized by the length of the protein. (B) Torsion of a protein normalized by the length of the protein. Both the normalized Writhe and normalized Torsion of the proteins in the PDB ensemble show a bimodal distribution with a peak at a positive and a negative value, skewed to the right. However, the two distributions are different, indicating that the two parameters capture different aspects of the protein conformation.

The normalized Torsion values are shown in Figure 10(B). We point out that the distributions of Writhe and Torsion are apparently different, which may be expected, since the two parameters capture different characteristics of the 3-dimensional conformation of the proteins. The distribution of the Torsion is more complex, possibly bimodal as well, with two peaks at positive and negative values of Torsion, respectively (the negative peak being weak). Similarly to the Writhe, the Torsion values may be affected by the secondary structure elements.

It is interesting to point out that the pattern of peaks at positive and negative values for the global distributions was also observed for the local Writhe and Torsion distributions.

